# Genomic analysis of family data reveals additional genetic effects on intelligence and personality

**DOI:** 10.1101/106203

**Authors:** W. David Hill, Ruben C. Arslan, Charley Xia, Michelle Luciano, Carmen Amador, Pau Navarro, Caroline Hayward, Reka Nagy, David J. Porteous, Andrew M. McIntosh, Ian J. Deary, Chris S. Haley, Lars Penke

## Abstract

Pedigree-based analyses of intelligence have reported that genetic differences account for 50-80% of the phenotypic variation. For personality traits these effects are smaller, with 34-48% of the variance being explained by genetic differences. However, molecular genetic studies using unrelated individuals typically report a heritability estimate of around 30% for intelligence and between 0% and 15% for personality variables. Pedigree-based estimates and molecular genetic estimates may differ because current genotyping platforms are poor at tagging causal variants, variants with low minor allele frequency, copy number variants, and structural variants. Using ∼20 000 individuals in the Generation Scotland family cohort genotyped for ∼700 000 single nucleotide polymorphisms (SNPs), we exploit the high levels of linkage disequilibrium (LD) found in members of the same family to quantify the total effect of genetic variants that are not tagged in GWASs of unrelated individuals. In our models, genetic variants in low LD with genotyped SNPs explain over half of the genetic variance in intelligence, education, and neuroticism. By capturing these additional genetic effects our models closely approximate the heritability estimates from twin studies for intelligence and education, but not for neuroticism and extraversion. We then replicated our finding using imputed molecular genetic data from unrelated individuals to show that ∼50% of differences in intelligence, and ∼40% of the differences in education, can be explained by genetic effects when a larger number of rare SNPs are included. From an evolutionary genetic perspective, a substantial contribution of rare genetic variants to individual differences in intelligence and education is consistent with mutation-selection balance.

---

The scores of different types of cognitive ability tests correlate positively and the variance that is shared between tests is termed general intelligence, general cognitive ability, or *g.*^1^ General intelligence typically accounts for around 40% of the overall variance among humans in batteries that contain tests of diverse cognitive abilities. The personality traits of extraversion and neuroticism are two of the five higher-order personality factors that are consistently identified in dimensional models of personality. High levels of extraversion are associated with positive affectivity and a tendency to engage with, and to enjoy, social situations. High levels of neuroticism are associated with stress sensitivity, as well as mental and physical disorders.^2^ All of these traits are partly heritable, but have also been linked to evolutionary fitness. This paradox, that cognitive ability and personality appear to be under selective pressure yet retain heritable variation, could be resolved if rare variants, which are less amenable to selection, are found to play a major role in the genetic contribution to variance in these traits. We test whether genetic variants not in linkage disequilibrium with genotyped single nucleotide polymorphisms (SNPs) (including rare variants, copy number variants, and structural variants) make a contribution to intelligence and personality differences using two separate methods.

Firstly, using a recently-developed analytic design for combined pedigree and genome-wide molecular genetic data, we test whether rare genetic variants, copy number variants (CNVs), and structural variants make an additional contribution to the genetic variance in intelligence, neuroticism, and extraversion. Secondly, using unrelated individuals, and genotype data imputed using the Haplotype Reference Consortium^3, 4^ (HRC) data, we use minor allele frequency (MAF) stratified GREML (GREML-MS) to quantify the effect of SNPs with a MAF of ≥0.001 to determine if this additional variance can also be recovered based on SNPs alone using imputation.

General intelligence has been found to be heritable, with twin and family studies estimating that 50% to 80%^5^ of phenotypic variance is due to additive genetic factors, a proportion that increases with age from childhood to adulthood.^6^ Heritability can also be estimated from molecular genetic data. Using the genomic-relatedness-matrix restricted maximum likelihood single component (GREML-SC) method, the additive effects of common SNPs are estimated to collectively explain between 20% and 50% of variation in general intelligence,^7, 8^ with an estimate of around 30% in the largest studies.^9^ General intelligence is also a significant predictor of fitness components including mortality,^10^ fertility,^11, 12^ and higher social status,^13^ as well as mental and physical disease.^6^ General intelligence is associated with developmental stability,^14, 15^ suggesting that it is not selectively neutral.

As directional selective pressure on a trait is expected to deplete its genetic variation, the existence of such robust heritability findings seems paradoxical when evolutionary theory is considered.^16^ However, mutation-selection balance provides an explanation of how genetic variation can be maintained for quantitative traits that are under directional selective pressure. Mutation-selection balance describes instances where mutations that are deleterious to the phenotype occur within a population at the same rate that they are removed through the effects of selective pressure. Due to the removal of variants with deleterious effects on the phenotype, the existence of common variants with medium to large effects is not expected under mutation-selection balance. This is consistent with the current findings from large GWAS on cognitive phenotypes, including general intelligence and education, where common SNPs collectively explain a substantial proportion of phenotypic variance, but the individual effect size of each genome-wide significant SNP discovered so far is around 0.02%.^17, 18^

Population genetic simulations show that very rare (MAF < 0.1%) variants explain little of the population variance in traits that are not under selection.^19^ However, the contribution made by rare variants increases when their effects on a trait and on fitness are correlated either through pleiotropy, or by the trait directly affecting fitness.^19^ The genetically informative evidence that is available tends to show that variants associated with intelligence are also linked to better health,^20, 21^ although these effects may be outweighed by a negative effect on fertility.^22, 23^ There is also evidence that the regions of the genome making the greatest contribution to intelligence differences have undergone purifying selection.^24^ Whereas this does not necessarily imply that intelligence has been selected for or against across our evolutionary history, it does indicate that genetic variants that are associated with intelligence are also associated with fitness, which suggests that rare genetic variants and hence mutation-selection balance, may act to maintain intelligence differences.^19^

Empirical studies so far have failed to find evidence of a link between intelligence and rare variants.^25^ These studies have often been limited in scope, with only copy number variants or exonic regions being considered, or being limited in statistical power because all rare variants were treated as having the same direction of effect through the use of burden tests.^25–29^ Where such tests have found an association these have been in small samples and subsequently failed to replicate.^30^ However, in large samples, rare variants found within regions of the genome under purifying selection have been found to be associated with educational success,^31^ an effect that was greater for genes expressed in the brain. Hence, rare variants found in some genes appear to have an effect on intelligence.

Less is known about the genetics of personality.^32^ As with intelligence, heritability estimates for extraversion and neuroticism are much higher, around 34-48%, when based on quantitative (twin- and family-based) genetic methods^33^ compared to molecular genetic estimates (4–15% for neuroticism^34^ and 0–18% for extraversion^35, 36^). Both extraversion and neuroticism are predictive of social and behavioural outcomes as well as anxiety, well-being and fertility.^37–40^ Positive genetic correlations have been reported for extraversion with attention deficit hyperactivity disorder and bipolar disorder, and for neuroticism with depression and anorexia nervosa.^36^

In the current study, we quantify the total genetic effect from across the genome on intelligence (including education, which shows strong genetic correlations with general intelligence^41^ and is used as a proxy phenotype for it in genetic studies^42^), extraversion, and neuroticism. Two recent approaches allow us to include genetic variants not normally captured using GWAS. Firstly, as our sample included nominally unrelated individuals with varying degrees of genetic similarity, as well as family members who all provided genome-wide SNP data, we were able to decompose two genetic sources of variance corresponding to genetic effects associated with common SNPs at the population level (h_g_^2^), and genetic effects associated with kinship (h_kin_^2^) (i.e. associated with SNPs on a family basis). Among related individuals, linkage disequilibrium is stronger and hence allows us to capture variation not tagged by common SNPs at the population level. This includes rare variants, CNVs, and other structural variants. As the inclusion of family members can introduce confounding between shared genetic effects and shared environmental effects,^43^ we use the GREML-KIN method by Xia and colleagues^44^ to simultaneously estimate three sources of environmental variance: sibling effects, spouse effects, and family effects. By using information from both nuclear family relationships and the many more distant pedigree relationships in the cohort we analyse, this novel approach allows us to estimate kin-specific genetic variation net of common environmental effects. Secondly, we validate these findings using unrelated individuals by using genotypes imputed using the HRC panel.^4^ By using GREML-MS to derive a heritability estimate we were able include rare SNPs (MAF 0.001–0.01) as well as partition the SNPs by MAF to determine the contribution made to trait variation by rare variants.

## Materials and Methods

### Samples

Data was used from the Generation Scotland: Scottish Family Health Study (GS:SFHS).^4, 45, 46^ A total of 24 090 individuals (N_male_ = 9 927, N_female_ = 14 163, Age_mean_ = 47.6) were sampled from Glasgow, Tayside, Ayrshire, Arran, and North-East Scotland of whom 23 919 donated blood or saliva for DNA extraction. These samples were collected, processed, and stored using standard procedures and managed through a laboratory information management system at the Wellcome Trust Clinical Research Facility Genetics Core, Edinburgh.^47^ The yield of DNA was measured with a PicoGreen and normalised to 50ng/μl prior to genotyping. Genotype data were generated using an Illumina Human OmniExpressExome -8- v1.0 DNA Analysis BeadChip and Infinium chemistry.^48^ We then used an identical quality control procedure as Xia et al.^44^ that included removing SNPs not on autosomes or with a minor allele frequency (MAF) of <0.05, a Hardy-Weinberg Equilibrium *P*-value <10^−6^, and a missingness of >5%. This left 519 729 common SNPs from 22 autosomes. Following quality control, a total of 20 032 genotyped individuals (N_female_ = 11 804) were retained; 18 293 of these individuals were a part of 6 578 nuclear or extended families.^49^ The mean age of the sample was 47.4 years (SD = 15.0, range 18 to 99 years). The degree of the relationships found in GS:SFHS as well as the size of each of the matrices can be found in Table 1.

**Table 1.** Degree of relatedness in the 20 032 GS:SFHS data and number of pairwise relationships.

| Matrix | Number of non-zero off-diagonal entries |
| --- | --- |
| GRM<sub>g</sub> | 200 630 496 |
| GRM<sub>kin</sub> | 41 174 |
| GRM<sub>Family</sub> | 20 115 |
| GRM<sub>Sibling</sub> | 1 767 |
| GRM<sub>Couple</sub> | 8 495 |
| Degree of relationship | Number of pairs |
| 1<sup>st</sup> degree | 18 320 |
| 2<sup>nd</sup> degree | 7 851 |
| 3<sup>rd</sup> degree | 4 129 |
| 4<sup>th</sup> degree | 3 950 |
| 5<sup>th</sup> degree | 11 032 |
| Unrelated individuals | 200 585 162 |

For the G matrix all off diagonal entries are non-zero.
The distance of the relationship is identified using SNP relatedness and according to approximate ranges of the expected pair-wise relatedness, $0.5^{i-0.5}$ to $0.5^{i+0.5}$ for $i^{\text{th}}$ degree relatives.
Unrelated individuals defined as more than 5<sup>th</sup> degree relatives $r \leq 0.022$ .

### Ethics

The Tayside Research Ethics Committee (reference 05/S1401/89) provided ethical approval for this study.

### Phenotypes

General intelligence (*g*), years in education (Education), neuroticism, and extraversion were examined using GREML-KIN, and GREML-MS. Four cognitive tests were used to derive general intelligence; the Mill Hill Vocabulary Scale ((MHVS) test re-test reliability over 2 years 0.9, split half reliability r = 0.9),^50, 51^ the Wechsler Digit Symbol Substitution Task (DST) (test re-test reliability r = .90),^52^ Wechsler Logical Memory which measures Verbal declarative memory (split half reliability, part 1 = 0.88, part two 0.79)^53^ and executive function (phonemic Verbal fluency, using letters C, F, L) (Cronbach's alpha = 0.83).^54^ The general factor of intelligence (*g*) was derived by extracting the first unrotated principal component from the four cognitive tests. This single component accounted for 42.3% of the variance in the total sample and each of the individual tests used demonstrated strong loadings on the first unrotated component (DST 0.58, Verbal Fluency 0.72, MHVS 0.67, and Verbal declarative memory 0.63). Education was calculated in the GS:SFHS as the years of full time formal education which was recoded into an ordinal scale from 0 to 10 (0: 0 years, 1: 1-4 years, 2: 5-9 years, 3: 10-11 years, 4: 12-13 years, 5: 14-15 years, 6: 16-17 years, 7: 18-19 years, 8: 20-21 years, 9: 22-23 years, 10: > 24 years of education). Education and general intelligence were positively correlated (*r* = 0.38, SE = 0.01, p < 2.20 × 10^−16^).

The other two measures examined were the personality traits of extraversion and neuroticism which were measured using the Eysenck Personality Questionnaire Revised Short Form, a self-report questionnaire requiring a yes or no response to 24 items.^55^ Both scales have reliabilities of Cronbach's alpha > 0.85.^55^

The effects of age, sex and population stratification were adjusted for using regression prior to fitting the models in GREML. Supplementary section Figure 1 shows the number of principal components used to control for population stratification for each of the phenotypes used.

**Figure 1.**
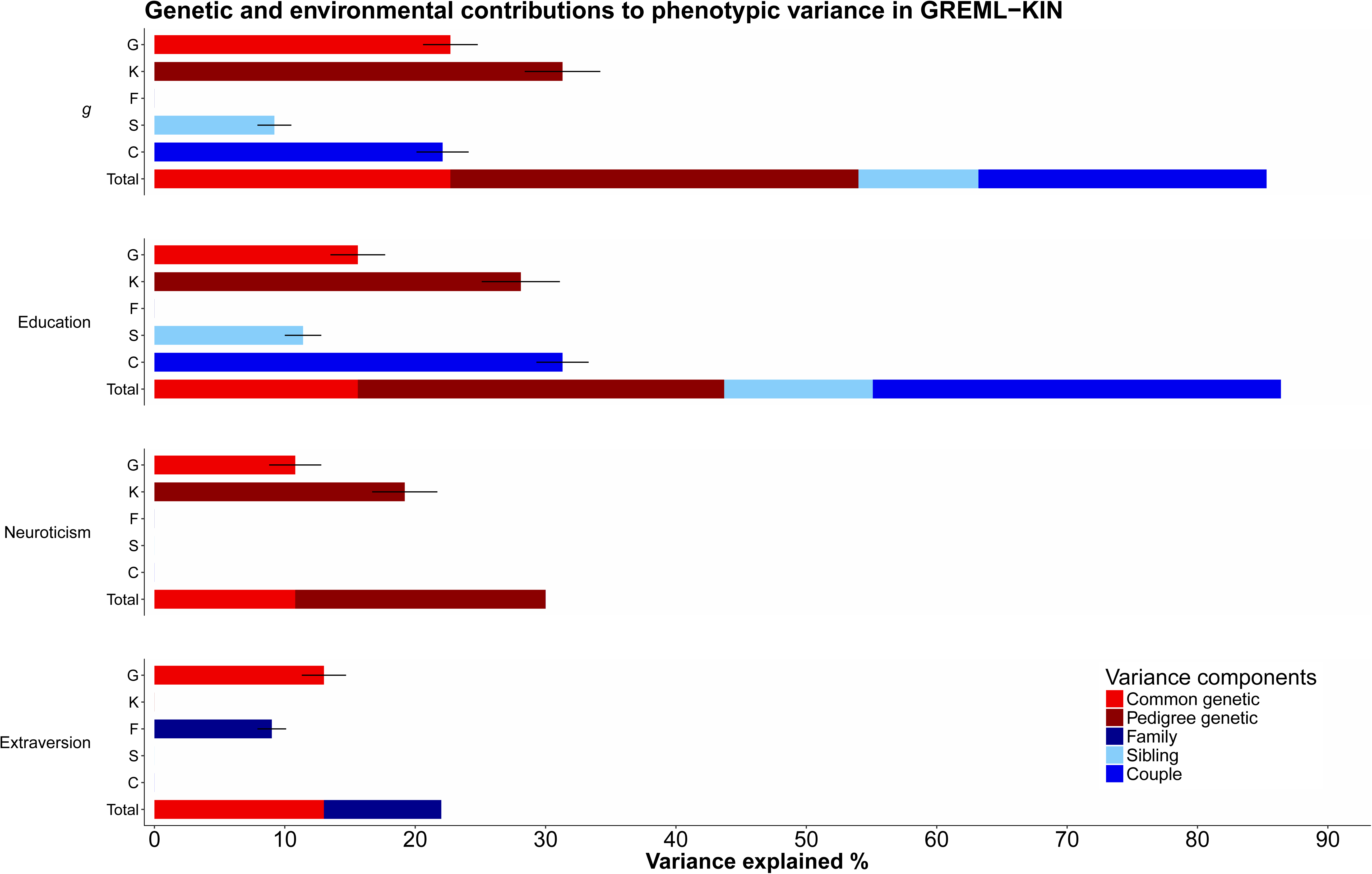
Selected models plotted for each of the phenotypes. Each component from the selected models is plotted individually, with the stacked bar plot showing the total proportion of the variance explained by the selected models. Error bars indicate standard errors.

### Statistical method

#### GREML-KIN: Partitioning phenotypic variance into five components

For each of the phenotypes examined here, variance was partitioned into five corresponding effects plus residual variance. This variance components analysis is based on the work of Zaitlen and colleagues^43^ who developed a method for estimating two genetic sources of variance in a data set with a measured family structure. Firstly, the variance component G can be estimated and used to derive h_g_^2^, the proportion of phenotypic variance explained by common SNPs, and secondly, the additional genetic effects associated with pedigree can be captured by K and used to derive h_kin_^2^, the proportion of phenotypic variance that is explained by genetic effects that are clustered within families. More recently this method has been extended by Xia and colleagues^44^ to include sibling, spouse, and nuclear family environmental effects. We refer to the extended method as GREML-KIN. The two genetic matrices described by Zaitlen et al. and Xia et al. model the effects associated with common SNPs (h_g_^2^) at the population level and those associated with pedigree (h_kin_^2^) respectively. These two genetic sources of variance were quantified using a genetic relationship matrix derived in the GCTA software.^56^

### Matrix construction

#### Genetic matrices

A genomic relationship matrix (GRM_g_) was used to derive the variance component of G in order to quantify the contribution made by common SNPs, h_g_^2^. This was derived in the manner set out by Yang and colleagues,^56^ where the estimated genomic relatedness between each pair of individuals is derived from identity by state SNP relationships and is found in each off diagonal entry in the GRM.). As the variance attributable to the shared environment was explicitly modelled here, no relationship cut off (typically, 0.025 is used) was applied to the genetic relationship matrix (GRM).

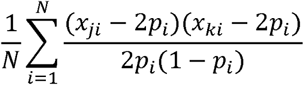

Minor allele frequency for SNP *i* is denoted as *p_i_* and the allelic dose (*x*) for individuals *j* or *k* at locus *i* is described as *x_ji_* or *x_ki_*. *N* indicates the total number of SNPs.

The kinship relationship matrix, GRM_kin_, (used to derive the variance component K) was derived using the method described by Zaitlen et al. (2013)^43^ by modifying the GRM_g_. Here, values in the GRM_g_ that were equal to or less than 0.025 were set to 0.

#### Environmental matrices

Three environmental matrices (ERM) were used to capture the variance associated with specific relationships between individuals. Each ERM was created by deriving an N by N matrix (where N is number of individuals) with diagonal entries set to 1 and non-diagonal entries set to 1 if the pair of individuals have the environmental relationship described or set to zero otherwise. The three ERMs derived here captured variance associated with the shared environment of spouses, (ERM_Couple_, variance component C), siblings (ERM_Sibling_, variance component S), and nuclear families (ERM_Family_, variance component F). As discussed by Xia et al.^44^ whilst these matrices are formed using information about the environment of an individual, they very likely will capture some effects of assortative mating (ERM_Couple_) as well as dominance effects (ERM_Sibling_), if any exist. Nevertheless, we retain the name of these matrices as described in the original paper by Xia et al.^44^ who was the first to include these matrices.

#### Estimating the phenotypic variance explained

For each trait we first fitted the two GRMs and the three ERMs simultaneously using a linear mixed model (LMM) using the GCTA software.^56, 57^ This full model is referred to as the GKFSC model, as it includes the genetic, kinship, family, sibling, and couple matrices.

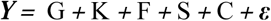

Here, ***Y*** is a vector of standardised residuals derived from one of the phenotypes. Random genetic effects were explained by fitting the G and K, which captured variants in LD with common SNPs found across a population and the extra genetic effects captured by the increase in LD found between members of the same extended family, respectively. Random environmental effects that were shared between related pairs of individuals were captured by fitting the F, S, and C to quantify the contributions made by environmental similarities between members of a nuclear family, siblings, and couples, respectively.

Restricted maximum likelihood (REML), implemented using the GCTA software,^56^ was used to estimate the variance explained by each of the variance components, with statistical significance determined using a log-likelihood ratio test (LRT) and the Wald test. Model selection began with the full GKFSC model (referred to as the full model). Components were dropped if they were not statistically significant according to both the Wald and the LRT tests. The model that contained only components that explained a significant proportion of variance is referred to as the selected model. If more than one component could be dropped from the model, we dropped the one with the worse fit first and then tested the significance of the other. The full results of each model can be seen in Supplementary Table 1.

The phenotypic variance explained by the variance components of G, K, S, F, and C used to derive h_g_^2^ (common SNP-associated effects), h_kin_^2^ (pedigree associated genetic effects), e_f_^2^ (shared family environment effect), e_s_^2^ (shared sibling environment effect), and e_c_^2^ (shared couple environment effect) were estimated (Table 2).

**Table 2.**
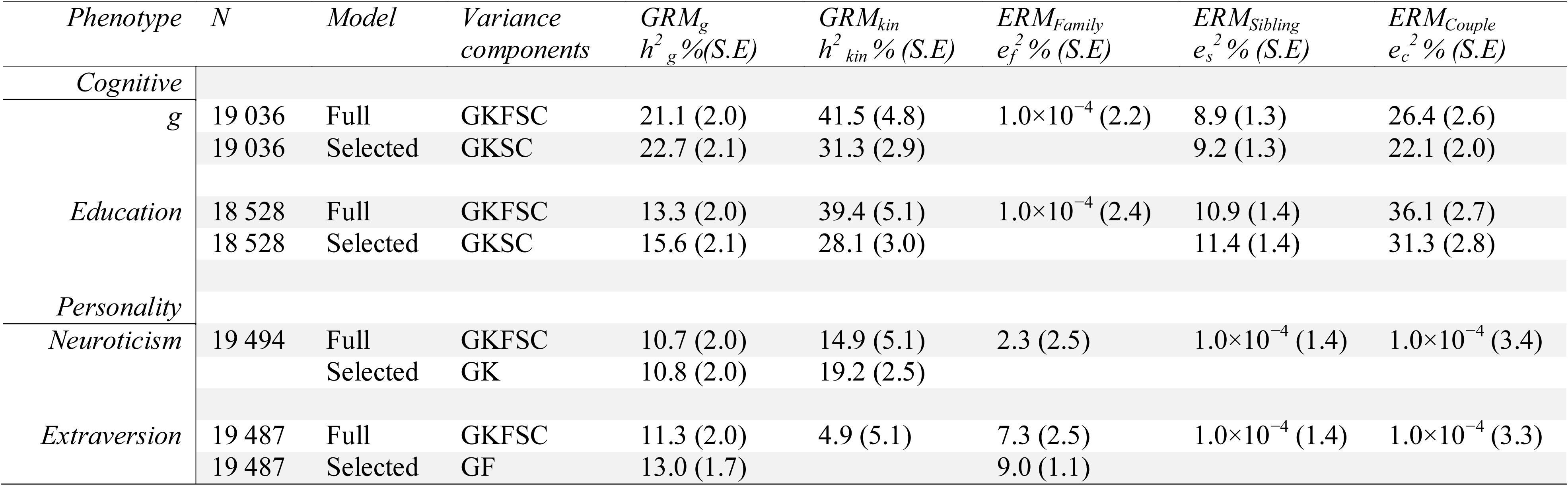
Results of variance components analyses for cognitive abilities and personality from the full model and the final model selected in a stepwise selection procedure.

Despite collinearity between the five matrices simulations conducted by Xia *et al*.^44^ show that this method provides robust results due to the dense relationships within the GS:SFHS cohort. The GS:SFHS is a family based cohort and the participants are related to varying degrees, including 1 767, 18 320, 7 851, 4 129, 3 950 and 11 032 pairs of couples, 1^st^, 2^nd^, 3^rd^, 4^th^ and 5^th^ degrees of relatives respectively. Therefore, what is shared between the ERM_Family_ matrix and GRM_kin_ matrix is information on the ∼18k 1^st^ degree relatives. However, ERM_Family_ holds ∼1.8k pairs of unique entries (couple pairs) and GRM_kin_ holds ∼23k pairs of unique entries (equivalent 2^nd^-5^th^ degree relative pairs of who were greater than 0.025 genetically identical). The unique entries from both matrices result in an increase of power, which allows the correct disentangling of the variance from those two different sources. An additional point is that the pedigree-associated genetic effects decay as the distance of the relationship increases, whereas nuclear family environmental effects do not. Thus, the fact that GS:SFHS consist of different classes of relatives, as well as the unique entries within the GRM_kin_ and ERM_Family_, helps to capture the property of pedigree-associated genetic variants. This logic extends to separating the variance from each of the environmental matrices. Although ERM_Couple_ and ERM_Sib_ are nested within the ERM_Family_, there are 9 853 pairs of unique entries (representing parents-offspring) within the ERM_Family_, which helps to separate the environmental effects. As shown by Xia *et al.*^44^, this method reliably identifies the major sources of variance that contribute to trait architecture. However, as with any method, effects become harder to detect as significant as they become smaller, since more power is needed for the reliable detection of small signals. This means that if one of the matrices only contributes to a small proportion of the overall phenotypic variance (e.g. less than 5% in GS:SFHS) its contributions will not be estimated reliably and the component will be dropped in the model selection procedure. However, any excluded minor component in the final model will have only a limited influence on the estimates of the major components that are retained in the final model. Thus, the major components we detected for each trait should be estimated reliably.

#### GREML-MS analysis

In order to show that the GRM_kin_ in GREML-KIN captures the contributions made by genetic variants poorly tagged by genotyped SNPs and is not confounded by the inclusion of close relatives we replicated our results using unrelated individuals. Using genotyped data imputed using the Haplotype Consortium (HRC)^3, 4^ data set allowed investigation into low frequency variants using the Sanger Imputation Service (https://imputation.sanger.ac.uk/). A quality control check was performed by checking autosomal haplotypes to ensure that strand orientation, reference allele, and position matched the reference panel. Data were then pre-phased using the Shapeit2 duohmm option provided by the Shapeit2 v2r837 software^58–60^, where the family structure of GS:SFHS was used to improve the imputation quality.^61^ Finally, an imputation quality score of info <0.4 was used to exclude poorly imputed variants and non-bi-allelic variants. This resulted in 11 497 491 bi-allelic SNPs with MAF > 0.001 available for analysis.

A relatedness cut off was applied to the participants of GS:SFHS of 0.025 resulting in a sample size of 7 370. Note, that relatedness was based on the GRM_g_, -i.e. estimated by using all genotyped common SNPs on the autosomes for the whole population. To show that the additional variance captured by our GRM_kin_ is due to less common variants, imputed and genotyped variants were assigned to one of six matrices describing the frequency of the minor allele. The six bins, and the matrices derived using them, were MAF = 0.001−0.01 (GRM_0.001−0.01_), MAF = > 0.01−0.1 (GRM_0.01−0.1_), MAF = > 0.1−0.2 (GRM_0.1−0.2_), MAF = > 0.2−0.3 (GRM_0.2−0.3_), > 0.3−0.4 (GRM_0.3−0.4_), MAF = > 0.4−0.5 (GRM_0.4−0.5_).^62^ These six matrices were then fitted simultaneously and analysed using REML.

## Results

The results of the full GKFSC models as well as the results of the selected models, can be seen in Table 2. For general intelligence (*g*) the final model was the GKSC model, allowing for a significant contribution from additive common genetic effects, additive pedigree-associated genetic variants, shared sibling environment, and a shared couple environment. For *g*, common SNPs (h_g_^2^) explained 23% (SE = 2%) of the phenotypic variation. Pedigree-associated genetic variants (h_kin_^2^) added an additional 31% (SE = 3%) to the genetic contributions to *g,* yielding a total contribution of genetic effects of 54% (SE = 3%) on *g*. The net contribution of modelled environmental factors to phenotypic variance in *g* was 31%. This was due to two sources of variance, shared sibling environment (e_s_^2^) and shared couple environment (e_c_^2^), that accounted for 9% (SE = 1%), and 22% (SE = 2%), respectively. As noted previously, these estimates could also include effects of dominance and assortative mating, respectively.

The GKSC model was also the selected model for education. As with general intelligence, pedigree-genetic variants accounted for the majority of the total genetic contribution to phenotypic variation in these traits. Pedigree-associated genetic variants explained 28.1% of the variation in education, whereas common SNP effects explained 15.6% (Figure 1.). The genetic results, i.e. SNP and pedigree contributions combined, for *g* and education are similar to the heritability estimates derived using the traditional pedigree study design in the same data set, which found a heritability estimate of 54% (SE = 2%) for *g* and 41% (SE = 2%) for education (Figure 2).^63^ This indicates that the genetic variants with the greater estimated cumulative effect on cognitive abilities are those that are poorly tagged on current genotyping platforms.

**Figure 2.**
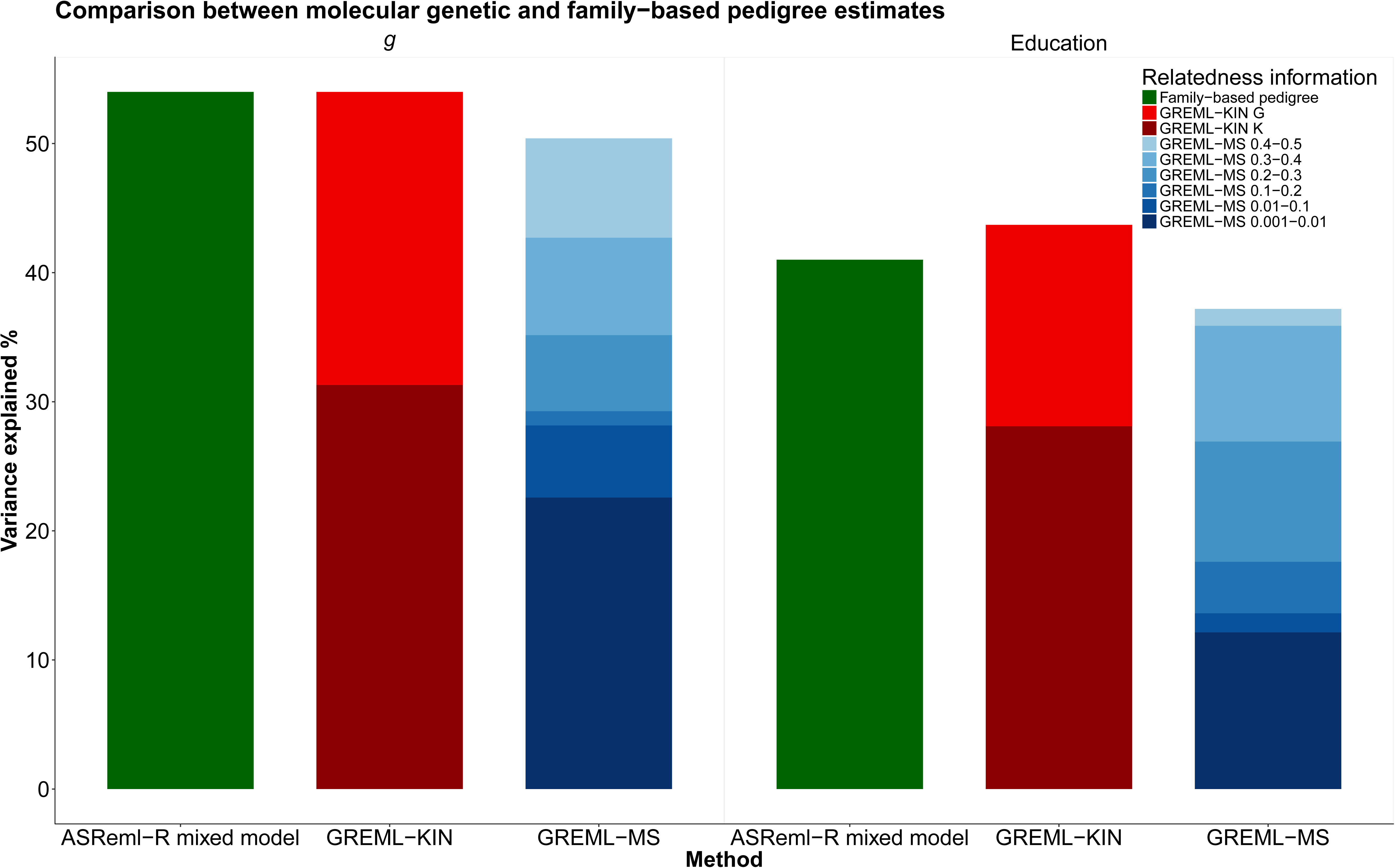
Bar plots showing the proportion of variance explained using family based methods and using molecular genetic data in related and unrelated samples. All of these analyses were performed using the same GS:SFHS data (*n* = 20 522, Education *n* = 22 406). Using related individuals and GREML-KIN, a sample size of 19 036 was available for general intelligence, and 18 5 280 for education after quality control. GREML-MS was conducted on unrelated individuals using a sample of *n* = 7 019 for general intelligence and 6 860 for Education. Estimates depicted in red were derived in the current study using GREML-KIN and show two sources of genetic variance. Bright red being common genetic effects captured by the GRM_g_ matrix and dark red being the additional genetic effects captured by exploiting the higher level of linkage disequilibrium between family members using the GRM_kin_ matrix. Estimates shown in shades of blue were derived using GREML-MS and indicate the variance explained using unrelated individuals with genotyped data imputed to the HRC reference panel. The estimates in dark green are taken from Marioni et al.^63^ and show the total genetic effects using ASReml-R mixed model when relatedness is inferred using identity by descent.

The results for each of the individual tests of cognitive ability used to derive general intelligence are each highly similar to general intelligence (Supplementary Table 2). For each of the single tests the K component captured a substantial and significant amount of phenotypic variance. The selected model for the Mill Hill Vocabulary test, the Verbal Fluency test, and Digit Symbol test was the GKSC model. The C component did not attain statistical significance for logical memory with the selected model being GKS.

For neuroticism the final model consisted of contributions from the variance components G and K. Additive common genetic effects explained 11% (SE = 2%) of the variance with pedigree-associated variants explaining an additional 19% (SE = 3%). Whereas none of the environmental components were statistically significant, the family component accounted for 2% of the variance in the full model and 1% in a model that included only the G and the K in addition to F.

For extraversion the only detectable source of genetic variation came from the G, which accounted for 13% (SE = 2%), with F explaining a further 9% (SE = 1%) of the phenotypic variation. The lack of pedigree-associated genetic effects could be due to low statistical power, as K explained 5% of the variance in the full model and 6% in a GKF model, but with a relatively large SE, estimated at 5%.

In addition to our model selection procedure, we also fit all possible component combinations for all phenotypes, to show a more complete account of the data and to give readers the ability to explore the consequences of including different components for the results, even when some of those components were not significant. The results have been made interactively available at https://rubenarslan.github.io/generation_scotland_pedigree_gcta/.

The results of GREML-MS are consistent with GREML-KIN. The total contribution of all SNPs resulted in a heritability estimate of 50% (SE = 10%) for intelligence and 37% (SE = 10%) for education (Table 3). This trend for the total heritability estimate derived from GREML-MS being similar to, but lower than, the heritability estimates derived from summing the G and K from GREML-KIN, and those derived from traditional pedigree-based methods (Figure 2) was evident across all cognitive variables. This attenuation is consistent with the findings of Evans et al.^64^ who showed that with imputation to HRC, GREML-MS can underestimate heritability by as much as 20% if the genetic architecture of a trait includes many rare variants.

**Table 3.** Results of GREML-MS variance components analyses for cognitive abilities and personality using six Minor Allele Frequency cut offs.

| *Phenotype* | *N* | *Minor allele frequency (MAF)* |  |  |  |  |  | Total variance explained<br>h<sup>2</sup> % (S.E) |
| --- | --- | --- | --- | --- | --- | --- | --- | --- |
|  |  | 0.001-0.01<br>h<sup>2</sup> % (S.E) | > 0.01-0.1<br>h<sup>2</sup> % (S.E) | > 0.1-0.02<br>h<sup>2</sup> % (S.E) | > 0.2-0.3<br>h<sup>2</sup> % (S.E) | > 0.3-0.4<br>h<sup>2</sup> % (S.E) | > 0.4-0.5<br>h<sup>2</sup> % (S.E) |  |
| *Number of SNPs* |  | 3 898 626 | 3 320 146 | 1 413 929 | 1 061 603 | 930 841 | 872 346 | 11 497 491 |
| *Cognitive* |  |  |  |  |  |  |  |  |
| *g* | 7 019 | 22.6 (9.5) | 5.6 (5.3) | 1.1 (3.5) | 5.9 (3.4) | 7.5 (3.3) | 7.7 (2.9) | 50.4 (9.9) |
| *Education* | 6 860 | 12.1 (9.6) | 1.5 (5.2) | 4.0 (3.6) | 9.3 (3.6) | 9.0 (3.4) | 1.3 2.8) | 37.2 (9.9) |
| *Personality* |  |  |  |  |  |  |  |  |
| *Neuroticism* | 7 195 | 1.0×10<sup>-4</sup> (8.8) | 3.6 (5.0) | 1.0×10<sup>-4</sup> (3.2) | 2.3 (2.9) | 0.9 (2.9) | 4.7 (2.6) | 11.4 (9.4) |
| *Extraversion* | 7 188 | 17.0 (9.2) | 1.0×10<sup>-4</sup> (4.7) | 1.0×10<sup>-4</sup> (3.2) | 1.1 (3.1) | 1.1 (3.0) | 1.8 (2.5) | 20.9 (9.6) |

When examining the variance explained by MAF using GREML-MS (Figure 3 and Figure 4) for general intelligence and education it is clear that the variants tagged by SNPs with a MAF between 0.001−0.01 make a large contribution to phenotypic variation. These low MAF variants explain 22.6% (SE = 9.5%) of the variation in intelligence, compared to 27.8% from variants with a MAF greater than 0.01. For education, low MAF variants explain 12.1% (SE = 9.6%), with all other variants explaining a total of 25.1%. Similar findings were also evident for each of the cognitive tests used in the general intelligence phenotype (Supplementary Table 3). This was also found for extraversion, where variants with a MAF of 0.001−0.01 explained 17.0% (SE = 9.2%) whilst all other SNPs explained only 4% of phenotypic variance. However, for neuroticism there was no evidence of any contribution made by the SNPs with a MAF of 0.001−0.01, and all variants only explained 11.4% (SE = 9.4%) of phenotypic variance.

**Figure 3.**
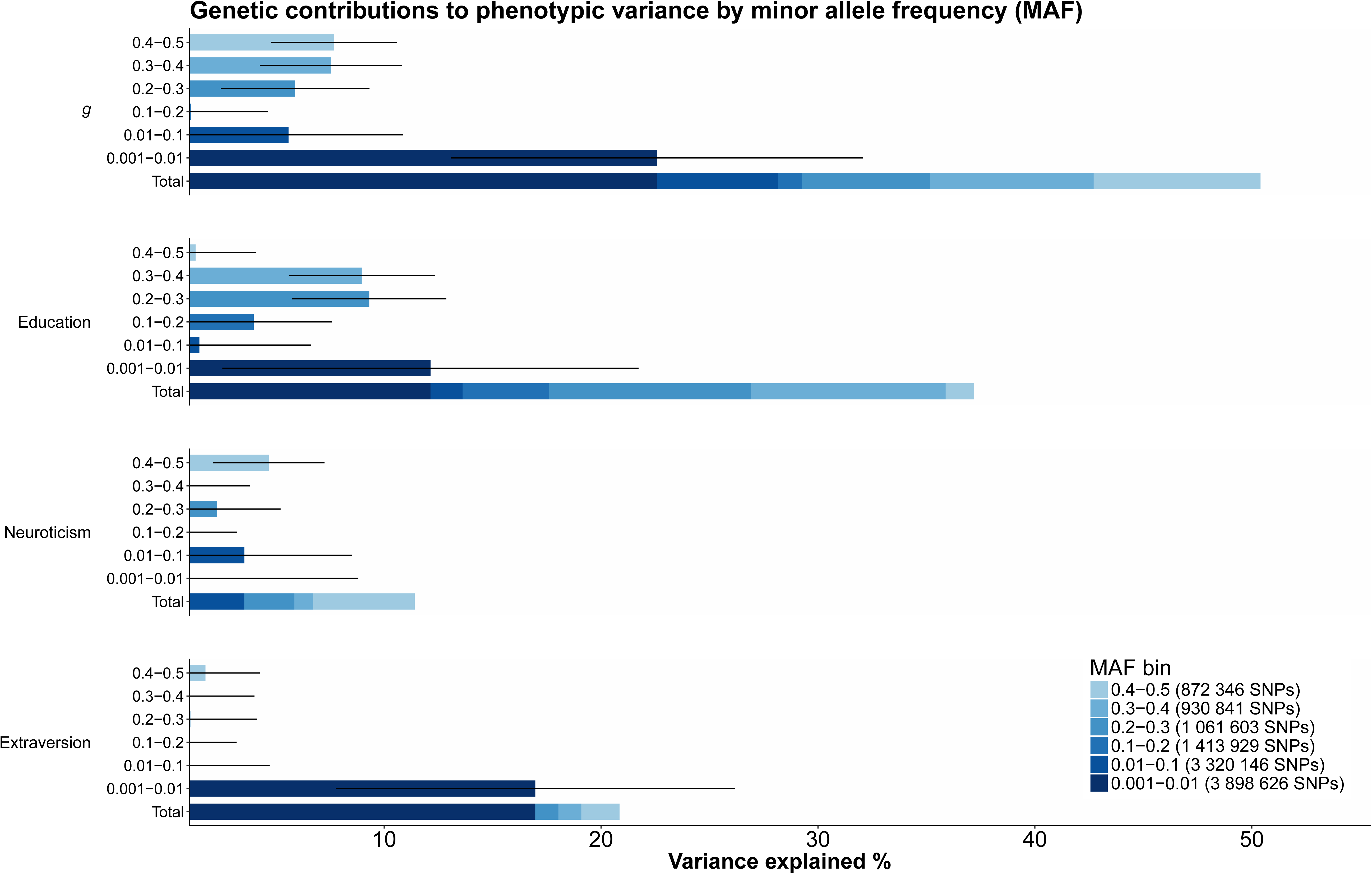
Genetic contributions to each of the phenotypes by MAF derived using GREML-MS. Each MAF cut off used is plotted separately, with the stacked bar plot showing the total proportion of the variance explained by the each MAF cut off. Error bars indicate standard error.

**Figure 4.**
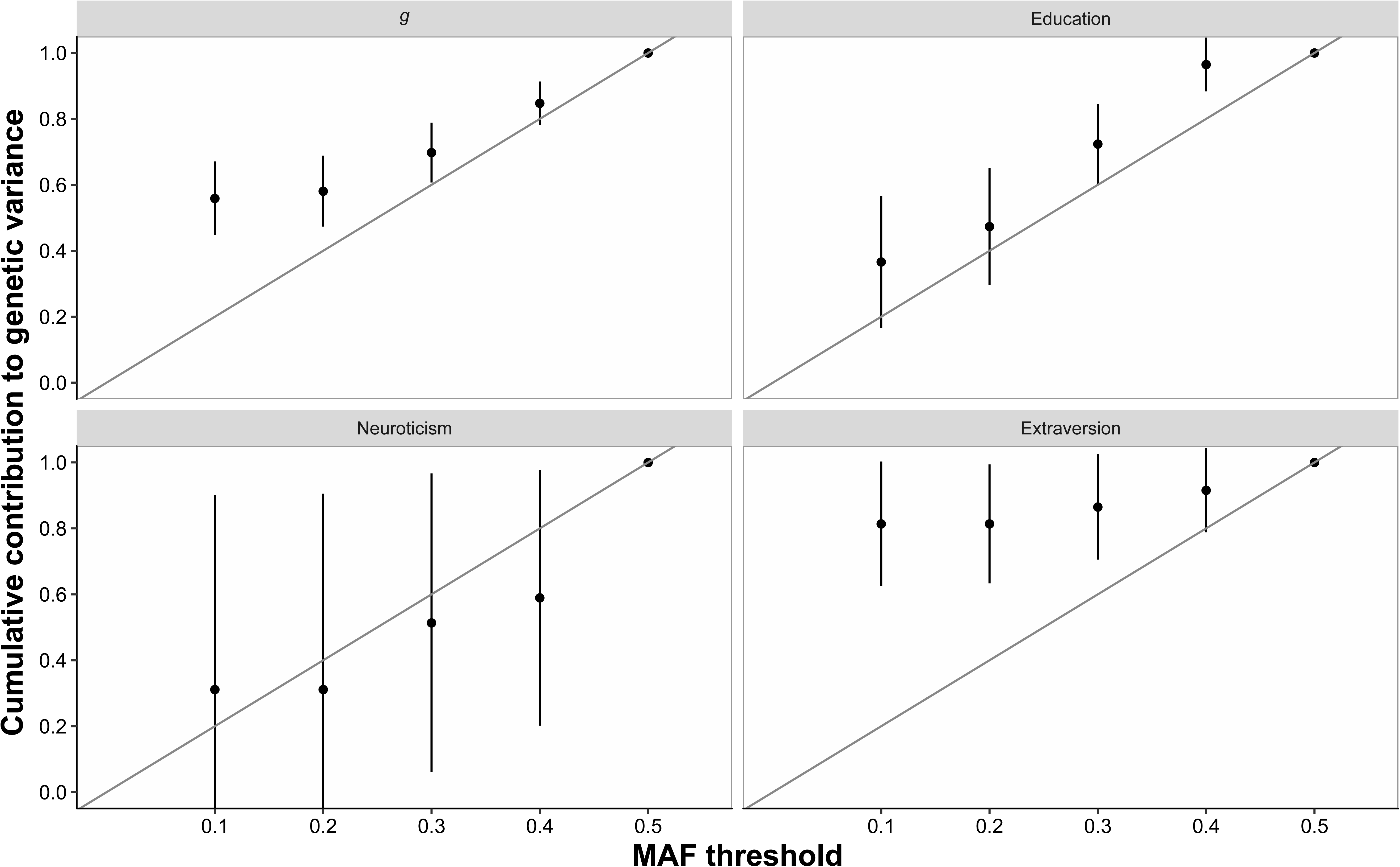
MAF plotted against the cumulative genetic variance explained. The diagonal grey line indicates evolutionary neutrality where the proportion of genetic variance is proportional to the MAF. Error bars represent standard errors for the cumulative variance components derived using the delta method, they are clipped if they leave the range of 0 to 1.^62^

We next examined if there was evidence of selective pressure acting on the cognitive and personality variables using GREML-MS. For a trait that is not under selective pressure whilst the majority of genetic variants will be rare, the majority of genetic variation associated with the trait is expected to be common.^65^ A trait that is evolutionary neutral will therefore show a linear proportional relationship between MAF and cumulative genetic variance explained.^62^ As can be seen in Figure 4 general intelligence shows a deviation from the neutral evolutionary model. Education, an often used proxy phenotype for intelligence^42^ showed no such deviation, indicating a different genetic architecture than that of general intelligence.

Extraversion also demonstrated evidence that low MAF variants made a greater contribution than more common variants. Neuroticism, however followed the model predicted under the assumption of evolutionary neutrality.

## Discussion

This study aimed to decompose and quantify genetic and environmental sources of variation to intelligence and personality in novel manners, using molecular genetic and pedigree data from the same large sample. In doing so, we sought to identify reasons for the gap between pedigree-based and SNP-based estimates of heritability in samples of unrelated individuals, a difference that might be due to genetic variants in poor linkage disequilibrium with SNPs genotyped on current platforms. A number of novel findings speak to long-standing questions in behaviour genetics and evolutionary genetics of psychological differences.^16, 32, 66^

Firstly, using GREML-KIN we could account for the entire heritability of general intelligence and education, as estimated in twin and family studies, by adding the G and K estimates we derived directly from genome-wide molecular genetic data.^63, 67^ Secondly, using GREML-MS, we replicated this finding with imputed data on unrelated individuals. For general intelligence and education, a substantial and significant proportion of the phenotypic variance was found to be explained by pedigree-associated genetic effects (h_kin_^2^). The pedigree-associated genetic variants accounted for over half of the genetic effects in these phenotypes. Even though GREML-MS is expected to underestimate heritability for traits where the genetic architecture includes the contribution of very rare variants,^64^ we were nevertheless able to recover the majority of this heritability following imputation to the Haplotype Reference Consortium. For neuroticism, G plus K estimates were ∼30%, even slightly exceeding the narrow-sense heritability estimates meta-analytically derived from family and adoption studies with heterogeneous measurements of personality.^33^ However, the K component was dropped for extraversion in our model selection procedure. Furthermore, results were less consistent between GREML-KIN and GREML-MS for personality traits. These convergences and divergences between both our methods and published results are potentially diagnostic for the genetic architecture of the traits under study.

The GREML-SC method of estimating heritability from unrelated individuals using common genome-wide SNPs often produces lower heritability estimates than those derived using family-based studies because it relies on LD between genotyped SNPs and causal variants at the population level. Should LD between genotyped SNPs and causal variants be low, then the genetic similarity between a pair of individuals at the causal variant will be different to the genetic similarity at genotyped SNPs, resulting in an underestimation of heritability. In within-family and twin studies, relatedness is based on identity by decent (IBD), where segments of DNA have been inherited from a recent common ancestor. Should a region be IBD between a pair of individuals, then all variants within that segment, except *de novo* mutations, are shared. Population-based SNP methods are sensitive to allele frequency, whereas IBD methods are blind to such effects. Therefore, the discrepancy between heritability estimates is consistent with the idea that causal variants in low LD with genotyped SNPs account for difference between IBD methods and population-based estimates derived using molecular genetic data.

In the current study we investigate if variants in poor LD with genotyped SNPs account for additional heritability by using DNA from close family members. Higher genetic relatedness within families leads to an increase in the LD between genotyped SNPs and potentially causal variants, resulting in heritability estimates in our study that are comparable to pedigree-based methods. This provides evidence that for intelligence the gap between the heritability estimates derived using IBD methods and those derived using SNP-based population methods is most likely due to causal variants in low LD with genotyped SNPs. In addition, we were able to model this missing variance and separate it from the additive common genetic effects that are estimated in a GREML-SC analysis based on unrelated individuals. The additional source of additive genetic variance from closely related family members, captured here in our kinship matrix (GRM_kin_), would be unmeasured in a GWAS on unrelated individuals using genotyped data. Whilst the use of related individuals can result in the confounding of pedigree genetic effects with shared family environmental effects, we were able to distinguish the contributions made to phenotypic variance by pedigree-associated genetic variants from those by shared family environment through modelling three sources of environmental variance,

The replication of the GREML-KIN findings with GREML-MS in the subsample of unrelated individuals provides strong evidence that the heritability estimates are not due to confounding with the environmental sources of variance modelled here. Indeed, both of these methods provide highly similar estimates, which are in turn similar to the estimate found using traditional pedigree-based analyses,^63^ indicating that the total heritability of intelligence can be captured using GREML-KIN. When using genotyped or imputed data, GREML-MS has been shown to underestimate the contribution made by rare variants to a polygenic traits by as much as 20%.^64^ This is most likely due to the low imputation quality of rare SNPs, which can be ameliorated by using whole-genome sequencing data (WGS) to derive a heritability estimate. However, for traits where very rare variants have an effect (minor allele count > 5), a downward bias is still apparent with WGS.^64^ GREML-KIN can also capture non-SNP associated variants like CNVs, which will also be missed by GREML-MS. This indicates that the accuracy of the heritability estimate provided by GREML-MS is dependent on the frequency of the causal variants that make up trait architecture, albeit much less so than using GREML-SC on genotyped data alone. Using GREML-KIN only a minor underestimation of heritability is seen, and regardless of MAF, heritability estimates are as accurate for genotyped data as they are for WGS. This suggests that, in the absence of environmental confounding, GREML-KIN approximates the true heritability better than GREML-MS. However, it should be noted that family-based analysis would be unsuitable for some phenotypes, such as those based on area or household measurements, as is the case with socioeconomic status or household income.^68^ Converging estimates from the different methods increase our confidence in their interpretation as genetic effects, whereas the divergences between methods can help diagnose potential unmeasured sources contributing to broad-sense heritability or confounding.

The patterns found in our GREML-MS analyses were consistent with the findings of Evans et al.^64^ for neuroticism and fluid intelligence. However, both GREML-KIN and GREML-MS estimates for neuroticism and extraversion fell short of estimates of broad-sense heritability in twin studies (47%^33^; 45%^69^). As previous research has suggested,^33, 70^ this is consistent with epistasis playing a major role in personality genetics, as a non-additive genetic component is not captured well outside of twin studies. Previous research^70^ did not discuss gene-environment correlation and interaction as a plausible cause for heritability estimates being higher in twin than in adoption and family studies, presumably because the shared environment contribution to personality variation was usually estimated not to be different from zero. Still, the difference between twin estimates of heritability and those presented here may also be explained to some extent by gene-by-environment interactions and gene-environment correlations.^32^

Another noteworthy divergence occurred between GREML-KIN and GREML-MS results for the personality traits. For extraversion, SNPs with a MAF of 0.001−0.01 explained 17.0% (SE = 9.2) whilst the K component explained only 4.9% (SE = 5.1) and was dropped from the final selected model. However, the G plus K estimate for extraversion is 17.9%, which is not significantly different from the total heritability estimate provided by GREML-MS (20.9%). This is consistent with the interpretation that there is an effect of the K component for extraversion which is too small to attain statistical significance in this sample. The results of neuroticism also do not match between GREML-KIN and GREML-MS. The total heritability estimate for GREML-MS was 11.4%, similar to the G estimate, but in GREML-KIN the K explained a further 19% (SE = 2.5), while almost no effect was found for SNPs with a MAF of 0.001−0.01 using GREML-MS. As the GREML-KIN estimate is closer to twin and family study estimates of the narrow-sense heritability for neuroticism, this discrepancy might mean that the causal variants involved in neuroticism are even rarer, or perhaps due to non-SNP-associated genetic variants captured by GREML-KIN, but missed in GREML-MS. Potentially, the slightly lower measurement reliabilities for our personality measures may explain why results are less consistent than for intelligence.

The pattern we found using GREML-KIN is consistent with rare variants explaining much of the gap between heritability estimates from pedigree and GREML-SC analyses, although CNVs and structural variation could also play a part, because they are poorly tagged by genotyped SNPs as well. This can be seen in Evans et al.^64^, who used two genomic matrices, corresponding to the GRM_g_ and the GRM_kin_ in the current study (for continuity, we will use our terms to describe their matrices). By varying the frequency of the causal variants in a simulated dataset, Evans et al. showed that even when using only array markers the total variance captured by these two matrices was equal to the true heritability in the data set, irrespective of the frequency of the causal variants. Consistent with the notion that the pedigree genetic effects captured by the GRM_kin_ are due to the effect of rare variants, GRM_kin_ captured an increasingly greater proportion of variance as the causal variant frequency fell. The reverse was true for the GRM_g_, which captured less variance as causal variant frequency fell.

We found further, more direct support for an important role of rare variants using GREML-MS, which showed that for each of the cognitive variables examined here, a large contribution to phenotypic variance was made by SNPs with a MAF between 0.001 and 0.01. For extraversion almost all of the heritability was tagged by low-MAF SNPs. Together this indicates that the genetic signal to be found in imputed GWAS is much larger than GREML-SC estimates based on genotyped unrelated individuals would suggest. For intelligence, almost all of the heritability can be found in GWAS with sufficient sample sizes, providing genotyped data are imputed using a high quality reference panel such as HRC.

In our GREML-MS results for general intelligence and extraversion the relationship between MAF and cumulative genetic variance explained was not proportionately linear, with increasing contributions being made to the genetic variance explained as MAF fell. This pattern contradicts the neutral evolutionary model^65^ and suggests that rarer variants have a larger effect on intelligence and extraversion. This is consistent with previous findings that genetic variance in regions of the genome that have undergone purifying selection also make the greatest contributions to intelligence differences.^24^

The GREML-KIN results favour the inclusion of a large K component for all traits except extraversion. This is consistent with a major contribution by rare and other poorly tagged variants. Previous work has already suggested role for mutation-selection balance acting on harm avoidance and novelty seeking,^71^ traits that are related to neuroticism and extraversion, respectively.^72^

Our variance analyses are blind to the direction of effects and the number of variants involved in each genetic component. If, as we would predict, future work finds that variants with the lowest minor allele frequencies tend to have larger negative effects on intelligence, it would imply a coupling between the selection coefficient of alleles and their effect on intelligence, as selective pressure would act to minimise the frequency of highly deleterious variants. If this coupling were strong,^73^ future work might infer that selection on intelligence was important in the past, even though current selective pressure goes in the opposite direction.^74^ If the impact of intelligence on fitness were limited to instances of pleiotropy with, for example, health, as some initial research suggests^20, 21^, the coupling between the selection coefficients of alleles and their effect sizes would be expected to be weaker. Selective pressure would act on the health-linked variants, whereas intelligence-linked variants would only be selected to the extent of their pleiotropic effects on health. This would de-couple the selection coefficient of an allele and its effect on intelligence. Therefore, such analyses could disentangle how much directly or indirectly intelligence has been under selection. Future work can use the SNPs known to affect intelligence and personality^17, 18^ to empirically quantify the coupling between allele frequency (indicating selection strength) and effect size in order to test this explanation directly, as has been demonstrated for height and BMI.^62^ Targeted re-sequencing of enriched genetic regions^24, 75, 76^ might be necessary to find very rare genetic variants associated with intelligence and personality, as has proven fruitful for example in prostate cancer research.^77^

The sibling component, which was retained in all models of intelligence, tracks the meta-analytic estimate of shared environmental variance (11%) from twin studies almost exactly. However, in our study the sibling component might also include the quarter of the dominance variation that siblings share, because siblings are the only relationship in this study where dominance plays a significant role.^44^ In the classical twin design, dominance variation (making dizygotic twins more dissimilar than half the similarity of monozygotic twins) can be obscured by shared environment effects (making dizygotic twins more similar). There is some evidence from other approaches that dominance only plays a minor role in intelligence differences.^78–81^

The family component was only retained in the model for Extraversion, although the point estimate was non-zero in the full Neuroticism model as well. This is consistent with meta-analytic estimates of shared environment for adults,^69^ and so may indicate a lack of power in the present study to detect these small effects. However, it may also be due to some level of confounding between K and F, where the association between extraversion and the F is due to contributions of the genetic factors accounted for by the K.

The couple component is somewhat complex to interpret. For intelligence and education, there is evidence of assortative mating,^82^ which will increase both the genetic and environmental similarity between couples. The couple component may mostly reflect this spousal similarity, along with the effects of more recent environmental influences. Beyond that, intelligence is not perfectly stable across the life course and studies of twins in earlier childhood frequently find a sizeable shared environment component. The importance of shared environment is usually said to decline from childhood to adulthood,^83^ as individuals pick their environmental individual niches (i.e., active gene-environment correlation), but this is based only on environment shared with siblings. However, it may also be that the *current* environment remains important and that the spouse is a better aggregated indicator of the current environment than the sibling with whom one usually no longer shares a home in adulthood. We find no couple component for personality, which is consistent with much weaker assortative mating on personality, especially neuroticism and extraversion.^84–86^

In the current study we were able to exploit the high LD found between members of the same family to estimate the contribution of genetic effects that are normally missed in GREML-SC analyses of GWAS data. Using GREML-KIN, we simultaneously modelled the effect of the family, sibling, and couple environment to avoid potential environmental confounds inflating our estimates. For intelligence and education, we find that genetic variants poorly tagged on current genotyping platforms explained a substantial proportion of the phenotypic variance, raising our heritability estimates to match those derived using pedigree-based quantitative methods. Such variants can include CNVs, structural variants, and rare variants. We find similar effects for neuroticism. For extraversion, pedigree-associated variants appear to play a smaller role in phenotypic variation. GREML-MS analyses, used with data imputed to the HRC reference panel, allowed us to examine lower frequency variants in a sample of unrelated individuals and provides strong convergent evidence, especially for intelligence and educational attainment. Taken together, our results suggest mutation-selection balance has maintained heritable variation in intelligence, and potentially to some degree also in neuroticism and extraversion, explaining why differences in these traits persist to this day despite selection. Future work should directly measure rare variants, as well as CNVs and structural variants, and test the direction of their effects.

